# Structural and functional signatures distinguish gain- and loss-of-function cancer mutations across diverse protein families

**DOI:** 10.64898/2026.11.26.769491

**Authors:** Biniam Tsegay Haile, Frances Mary Genevive Pearl

## Abstract

Gain-of-function (GOF) and loss-of-function (LOF) missense mutations are major drivers of cancer, with contrasting effects on protein function. Although computational predictors can distinguish pathogenic from benign variants fairly accurately, they generally do not identify the underlying mechanism of pathogenicity, limiting biological interpretation of cancer-associated missense mutations.

We assembled a curated dataset comprising 1,727 GOF and 2,857 cancer-associated missense mutations whose assignment was supported by clinical or experimental evidence and comprehensively annotated each variant using structural, functional, evolutionary and pathogenicity features. Analysis of a range of pathogenicity predictors suggesting that predictors trained predominantly on germline or evolutionary variation may underestimate the pathogenic potential of somatic cancer mutations. Comparative analyses also demonstrated that GOF and LOF mutations tend to occupy distinct structural and functional environments in the proteins they affect. GOF mutations are enriched within regulatory domains, protein–protein interaction interfaces and phosphorylation-associated regions, while largely preserving overall protein structure, whereas LOF mutations often disrupt structurally constrained protein cores through destabilising substitutions. Guided by these observations, we developed biologically informed composite features that capture higher-order relationships between complementary protein descriptors. An XGBoost classifier, MOKCa-3D-GL, achieved a cross-validation AUROC of 0.939 and an independent validation AUROC of 0.834 for GOF versus LOF classification. The curated annotations generated in this study have been incorporated into the MOKCa-3D database, providing a benchmark resource for future computational method development.

Cancer-associated GOF and LOF missense mutations tend to exhibit distinct structural and functional signatures that can be exploited for accurate mechanistic prediction. By integrating interpretable structural and functional descriptors with biologically informed feature engineering, we provide both a predictive framework and a curated community resource that advance functional interpretation of cancer-associated missense mutations.

**Author summary:** Cancer is often driven by mutations that alter the way proteins’ function. Some mutations increase protein activity or cause proteins to behave abnormally (gain-of-function), while others reduce or eliminate their normal activity (loss-of-function). Although existing computational tools can predict whether a mutation is harmful, they often cannot explain how it affects protein function. Understanding these differences is important for uncovering how mutations contribute to cancer development.

In this study, we analysed more than 4,500 cancer-associated mutations to identify characteristics that distinguish these two mechanisms. We found that gain-of-function mutations tend to occur in regions that control protein activity or interactions with other proteins, while largely preserving their overall structure. In contrast, loss-of-function mutations frequently affect regions essential for maintaining protein stability and structure. We also found that some existing prediction tools may underestimate the harmful effects of mutations acquired during cancer development.

Using these findings, we developed a computational tool, MOKCa-3D-GL, that distinguishes between gain-of-function and loss-of-function mutations. Our curated data is available through the MOKCa-3D database. These resources provide new opportunities to understand how individual mutations contribute to cancer and may help guide future investigations into cancer biology and therapeutic strategies.

## 1. INTRODUCTION

Cancer develops through the progressive accumulation of somatic mutations that confer a selective advantage to tumour cells [1, 2]. Although large-scale cancer genome sequencing has transformed our understanding of the mutational landscape of human tumours, interpreting the functional consequences of individual missense variants remains a major challenge [3]. While only a minority of somatic variants function as cancer drivers, distinguishing these from biologically neutral, passenger mutations is essential for understanding tumour biology and informing precision oncology [4, 5].

Driver missense mutations alter protein function through distinct molecular mechanisms. Gain-of-function (GOF) mutations activate oncogenes by enhancing or altering protein activity, frequently through recurrent substitutions at key positions in the protein structure [6, 7]. In contrast, loss-of-function (LOF) mutations typically inactivate tumour suppressors through structural destabilisation, disruption of catalytic residues or loss of critical molecular interactions [7]. Although both mutation classes are pathogenic, the functional consequences arise through fundamentally different structural and biochemical mechanisms, suggesting that they should be distinguishable using protein-level features.

Numerous computational methods have been developed to predict the functional impact, deleteriousness or pathogenicity of missense variants, including PolyPhen-2 [8], SIFT [9], Mutation Assessor [10], FATHMM [11], EVE [12], ESM [13], and AlphaMissense [14]. However, recognition that somatic cancer mutations differ from inherited disease variants led to the development of cancer-specific predictors such as CHASM [15] and FATHMM-Cancer [11],which distinguish driver from passenger mutations.

Complementary structure-based approaches, including the Missense3D suite of predictors and databases [16, 17], together with integrative annotation platforms such as ProtVar [18], provide structural and functional context to support interpretation of missense variants. However, these approaches are primarily designed to assess variant impact or structural consequences rather than distinguish the molecular mechanisms by which mutations alter protein function.

More recently, machine learning approaches have extended prediction beyond pathogenicity to infer molecular mechanism. LoGoFunc predicts GOF, LOF and neutral variants using protein-, gene- and variant-level features [19], while VPatho applies deep learning to infer molecular mechanism [20]. Both methods incorporate multiple classes of genetic variation, including nonsense, frameshift and splice-site mutations, whose functional consequences are often more readily inferred than those of missense variants. Using predominantly germline disease mutations, Gerasimavicius et al. [21] showed that GOF, LOF and dominant-negative variants exhibit distinct structural signatures, with GOF mutations generally causing smaller structural perturbations than LOF mutations. Recently, Singh et al, 2025 demonstrated that activating and inactivating missense variants can be distinguished using structural and evolutionary features within protein kinases [22]. Cancer-specific resources have substantially improved the interpretation of somatic variants. COSMIC provides a comprehensive catalogue of somatic mutations [23, 24], OncoKB curates the biological effects, oncogenicity and clinical actionability of cancer variants [25, 26], and CIViC provides community-curated evidence describing the functional, oncogenic and clinical significance of cancer-associated variants [27]. MOKCa and the recently developed MOKCa-3D focus on documented gain-of-function and loss-of-function annotations integrated with structural and functional protein features [28, 29].

Experimentally documented GOF and LOF cancer missense mutations remain relatively scarce, and mechanistic annotations are fragmented across individual studies and knowledgebases, limiting the development of robust supervised models for mechanistic prediction.

Here, we present a structure-informed framework for mechanistic classification of cancer-associated missense mutations. Using a manually curated collection of experimentally or clinically validated GOF and LOF cancer associated missense mutations, we 1) identify structural and functional features that distinguish between the functional consequences of mutations and develop interpretable machine-learning classifiers, and 2) provide a curated benchmark resource through the MOKCa-3D database.

## RESULTS

### Construction of a curated mechanistic mutation dataset

We manually assembled a curated dataset of 1,727 GOF missenses mutations from 200 proteins and 2,857 LOF missense mutations from 336 proteins from the literature, representing one of the largest collections of mechanistically annotated cancer missense mutations currently available. We then extracted 3,831 non-pathogenic missense variants from 334 proteins in ClinVar [30]and termed these mutations as “neutral” as they had been explicitly not linked to any disease processes including cancers. Each variant was comprehensively annotated using complementary structural, functional, evolutionary and sequence-derived features, including residue conservation, solvent accessibility, structural environment, protein–protein interaction interfaces, post-translational modification sites, pathogenicity scores and AlphaFold confidence metrics (Fig S1).

The resulting dataset spans a diverse range of oncogenes and tumour suppressors with distinct structural architectures and molecular functions, providing a robust foundation for identifying general principles that distinguish GOF and LOF mutations across heterogeneous cancer-associated proteins.

### Distinct amino acid substitution signatures distinguish GOF and LOF mutations

GOF and LOF cancer mutations and ClinVar non-pathogenic variants exhibited distinct amino acid substitution patterns, indicating that different mutation classes are associated with characteristic physicochemical classes of amino acid exchange (Fig 1a, Supplementary Table S1).

**Figure 1.**
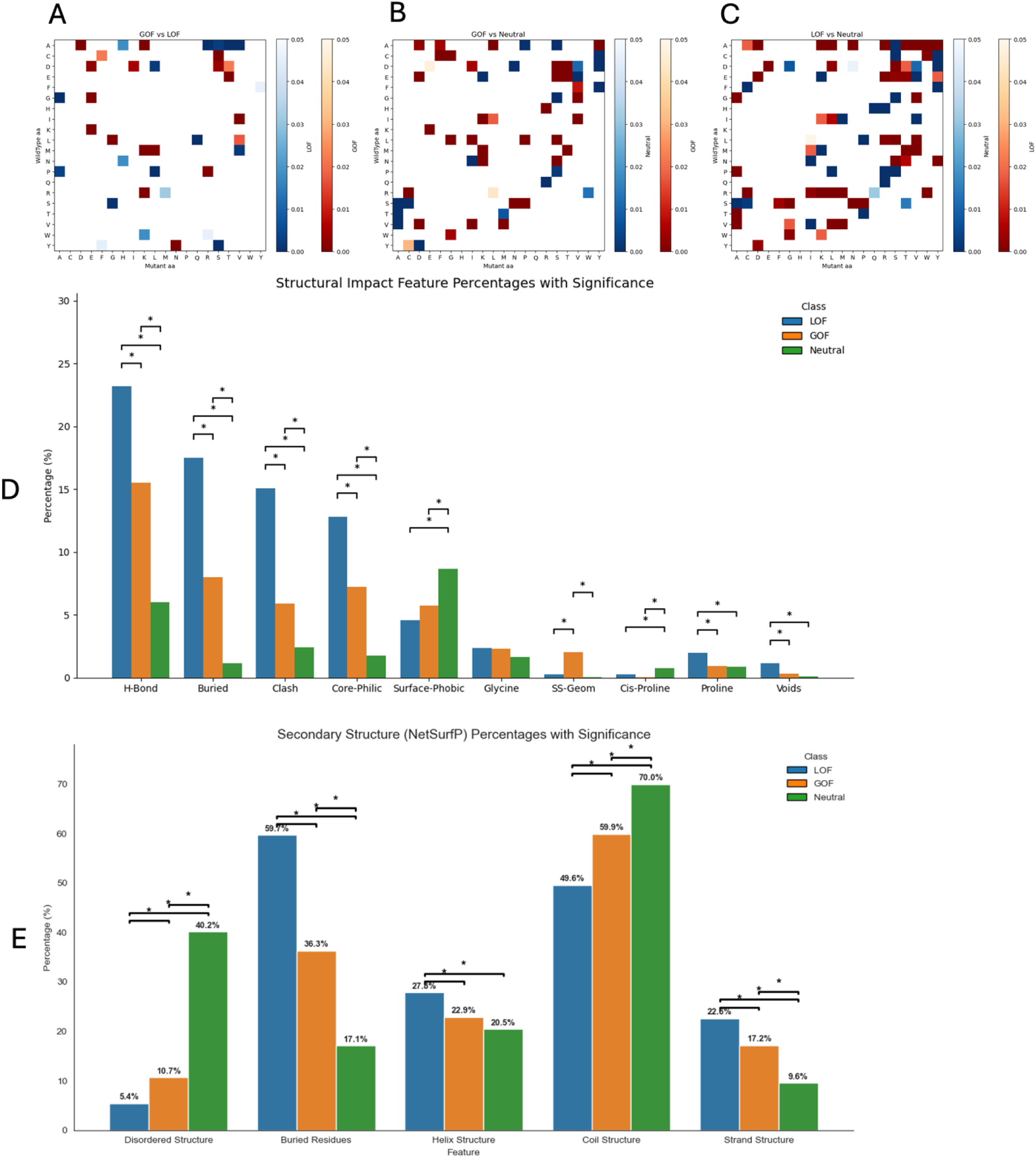
Structural and substitution characteristics of missense variants. (Across all panels, comparisons are made between gain-of-function (GOF), loss-of-function (LOF), and ClinVar non-pathogenic variants. Statistical significance (P < 0.05) was assessed using Fisher’s exact test with Bonferroni correction and is indicated by brackets or shading). Amino acid substitution enrichments: Heatmaps comparing mutation frequencies between classes A) GOF vs. LOF, B) GOF vs. ClinVar non-pathogenic and C) LOF vs. ClinVar non-pathogenic. Substitutions are mapped by wild-type (y-axis) and mutant (x-axis) amino acids. Darker shades indicate statistically significant enrichment (smaller P-values) for a substitution in one class over another. (D) Predicted structural perturbations: Bar plots showing the percentage of variants predicted by the SAAP-MOKCa-3D pipeline to introduce destabilizing effects. These include hydrogen bond disruptions, introducing a buried charge, steric clashes, hydrophilic core (CorePhilic), or hydrophobic surface (SurfacePhobic) substitutions. Mutations to/from glycine, disrupting a disulfide bond (SSGeom), mutations to/from proline and cavity formation (voids). (E) Structural properties of mutation sites: Percentage of variants in specific structural sites predicted by NetSurfP3 (coils, α-helices, β-strands, buried residues, and intrinsically disordered regions). Buried residues are defined as having relative solvent accessibility (RSA) ≤25%, and disordered residues are those with a probability >0.5.

GOF mutations displayed a relatively restricted substitution repertoire compared with LOF variants. Substitutions often involved changes in residue charge or polarity, with an enrichment of substitutions involving acidic residues (e.g. D→S, D→T and E→T). These patterns are consistent with local biochemical perturbations that may modify protein activity or molecular interactions without requiring extensive structural disruption. Recurrent substitutions introducing serine, threonine or lysine residues further suggest alterations to local regulatory chemistry, although the functional consequences of individual substitutions depend on their structural and sequence context.

In contrast, LOF mutations exhibited a broader spectrum of amino acid substitutions associated with larger physicochemical changes (See Fig 1b, Supplementary Table S1). The most common wild-type residues affected by LOF mutations were alanine and arginine. Enriched substitutions frequently involved substantial alterations in residue size, charge and hydrophobicity, including bulky aromatic and conformationally restrictive residues that are more likely to destabilise protein structure or disrupt native molecular interactions. ClinVar non-pathogenic variants (Fig 1c), by comparison, were enriched for conservative substitutions between chemically similar amino acids, consistent with preservation of normal protein structure and function.

Overall, the amino acid substitution profiles support distinct differences between GOF and LOF mutations. Whereas GOF mutations involved substitutions that are compatible with modification of protein function, LOF mutations more frequently introduced disruptive physicochemical changes associated with protein destabilisation.

### 2. 3 GOF and LOF mutations occupy distinct structural environments

To investigate the structural basis of mutation mechanism, we analysed residue-level structural descriptors generated by the SAAP-MOKCa-3D pipeline which calculates structural differences between the wild type and missense mutations, together with secondary structure, solvent accessibility and disorder predictions of the wild-type residue from NetSurfP3. Structural profiles differed most markedly between LOF and ClinVar non-pathogenic variants, while GOF mutations generally exhibited intermediate or feature-specific patterns (Fig 1).

GOF mutations occurred more frequently at solvent-accessible sites than LOF mutations (63.7% versus 40.3%), although the sites of ClinVar non-pathogenic variants remained the most solvent exposed (82.9%), consistent with their greater tolerance to amino acid substitution. Compared with LOF mutations, GOF variants showed substantially lower frequencies of features associated with protein destabilisation, supporting a mechanism in which protein activity is altered while overall structural integrity is largely preserved. The principal exception was disruption of disulphide bonds, which occurred in 2.03% of GOF mutations compared with 0.28% of LOF and 0.05% of ClinVar non-pathogenic variants: This enrichment may reflect mechanisms in which local structural rearrangements promote activation rather than global protein destabilisation, as exemplified by oncogenic RET C634R, where disruption of normal intramolecular disulphide bonding promotes aberrant disulphide-linked receptor dimerization and constitutive kinase activation [31].

In contrast, LOF mutations were consistently enriched for features associated with protein destabilisation (Fig 1d). Buried charge substitutions (17.5% versus 8.0% for GOF and 1.2% for ClinVar non-pathogenic variants), steric clashes (15.1% versus 5.9% and 2.4%), hydrogen bond disruption (23.2% versus 15.5% and 6.0%), and hydrophilic substitutions within the protein core (12.8% versus 7.2% and 1.8%) all occurred significantly more frequently in LOF mutations. NetSurfP3 analyses provided complementary evidence, showing that LOF mutations preferentially occurred at buried sites (59.7% versus 36.3% for GOF and 17.1% for ClinVar non-pathogenic variants) and were enriched within α-helices and β-strands, indicating that they predominantly disrupt ordered, structurally constrained regions in which amino acid substitutions are most likely to compromise protein stability and function (Fig 1e).

ClinVar non-pathogenic variants displayed a markedly different structural profile, being enriched within intrinsically disordered regions (40.2% versus 10.7% for GOF and 5.4% for LOF) and coil structures (70.0% versus 59.9% and 49.6%, respectively). Together with their enrichment for conservative amino acid substitutions, these observations are consistent with the greater mutational tolerance of structurally flexible protein regions.

Overall, these analyses demonstrate that GOF and LOF mutations often prefer distinct structural environments, with GOF mutations situated in solvent-accessible regions while largely preserving overall protein architecture, whereas LOF mutations predominantly disrupt structurally constrained protein cores. These complementary structural signatures provided the rationale for integrating structural descriptors with functional protein annotations to improve mechanistic classification.

### 2.3 Functional annotations reveal distinct mechanisms of pathogenic mutations

To investigate whether functional annotations distinguish the functional consequences of cancer-associated mutations, we compared prediction from the evolution-base models EVE [12], a LLM-based method ESM-1b [13] and conservation (ScoreCons) [32] obtained through ProtVar [18], with AlphaMissense [14] pathogenicity predictions, protein domain, post-translational modification (PTM) and protein–protein interaction (PPI) annotations across GOF, LOF and ClinVar non-pathogenic variants.

Overall, conservation and functional prediction methods clearly distinguished pathogenic cancer-associated variants from ClinVar non-pathogenic variants, whereas none clearly discriminated between GOF and LOF variants (see Fig 2, and Supplementary Fig S1) Among the predictors examined, AlphaMissense showed the strongest separation, classifying 76.0% of GOF and 85.4% of LOF mutations as pathogenic, while 82.4% of ClinVar non-pathogenic variants were classified as benign (Fig. 2d). These findings indicate that evolutionary conservation and pathogenicity prediction are effective at identifying functionally important cancer-associated variants but provide limited information about the direction of their functional effects. Additional structural and mechanistic features are therefore required to distinguish gain-of-function from loss-of-function effects.

**Figure 2.**
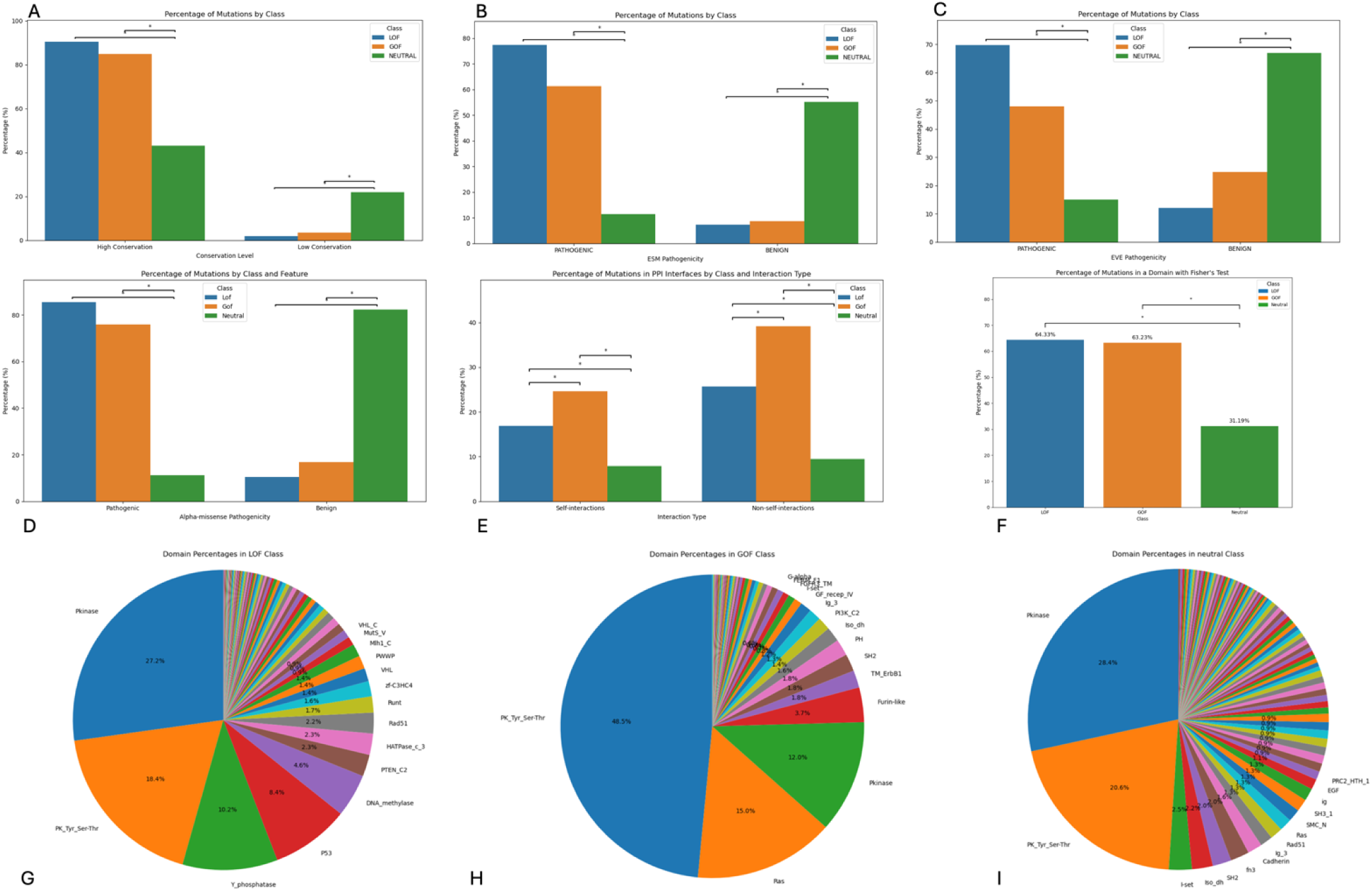
Distribution and characteristics of cancer-associated somatic mutations. (Across all panels, bar plots show the percentage of Loss-of-Function (LOF), Gain-of-Function (GOF), and ClinVar non-pathogenic mutations. Brackets with asterisks denote statistically significant pairwise differences using Fisher’s exact test). The percentages do not total 100% within each class; some variants were unclassified, ambiguous, or lacked scores. (A) Residue Conservation: showing percentage of mutation in High conserved and Low residues across mutation class). (B) ESM: Percentage of pathogenic versus benign mutation, determined by ESM pathogenicity score. (C) EVE: Percentage of pathogenic versus benign mutation, determined by EVE pathogenicity score. (D) Pathogenicity: Percentage of pathogenic versus benign mutations, determined by AlphaMissense pathogenicity scores. (E) Protein-Protein-Interaction (PPI) residues: Mutation frequencies at self-interaction and non-self-interaction PPI interfaces. (F) Protein domains: Proportion of mutations located within defined protein domains. (G)Pie chart to show Frequency of mutation in different protein domain of LOF variants dataset. (H) Pie chart to show Frequency of mutation in different protein domain of GOF variants dataset. (I) Pie chart to show Frequency of mutation in different protein domain of ClinVar non-pathogenic datasets. 90.48% of LOF and 84.94% of GOF variants occurred at highly conserved residues, compared with 43.15% of ClinVar non-pathogenic variants; both differences were statistically significant. Low-conservation scores were rare across LOF 1.96% and GOF 3.47% classes compared to ClinVar non-pathogenic 21.95% (Fig. 2a). ESM-1b classified 77.46% of LOF, 61.29% of GOF, and 11.47% of ClinVar non-pathogenic variants as pathogenic. In contrast, 55.26% of ClinVar non-pathogenic variants were classified as benign, compared with only 7.25% of LOF and 8.75% of GOF variants, with both comparisons reaching statistical significance (Fig. 2b). EVE produced clearer separation between functional and ClinVar non-pathogenic variants: 69.75% of LOF and 48.03% of GOF variants were classified as pathogenic, compared with 14.94% of ClinVar non-pathogenic variants, with 66.97% of ClinVar non-pathogenic variants classified as benign (Fig. 2c).

### Driver mutations preferentially target conserved functional regions

Both GOF (63.2%) and LOF (64.3%) mutations were significantly enriched within annotated protein domains compared with ClinVar non-pathogenic variants (31.2%) (Fig 2f), whereas no significant difference was observed between the two pathogenic classes. Protein kinase domains represented the most affected functional domain class, consistent with the central role of protein kinases in oncogenic signalling (Fig 2g-2i). This distribution may also reflect the enrichment of well-characterised kinase mutations in manually curated cancer mutation databases.

Despite the overall similarity in the frequency with which domains were affected, several protein domains showed distinct mechanistic preferences. GOF mutations were enriched within Ras family domains (15.0%), encompassing KRAS, NRAS and HRAS, together with furin-like cysteine-rich domains (3.7%) (Fig 2h) which mediate ligand recognition and receptor activation in extracellular signalling proteins. For example, glioblastoma-associated C311R and C334S substitutions in the extracellular furin-like cysteine-rich domain of HER2 are proposed to disrupt native intramolecular disulfide bonds, leaving unpaired cysteines that promote covalent receptor dimerization and oncogenic activation without directly altering the kinase domain [33].

Conversely, LOF mutations showed enrichment within protein tyrosine phosphatase domains (10.4%), p53 DNA-binding domains (8.4%), and DNA methyltransferase domains (4.6%) (Fig 2g). These domains are characteristic of tumour suppressor proteins, including PTEN, TP53 and DNMT3A, where loss of catalytic activity or DNA-binding capacity results in impaired regulation of cell signalling, genome integrity and epigenetic control. The preferential enrichment of LOF mutations within these domains is therefore consistent with established mechanisms of tumour suppressor inactivation.

### GOF mutations preferentially perturb protein-protein interfaces

GOF mutations were significantly enriched at protein–protein interaction interfaces. 39.1% of GOF mutations occurred at heteromeric interfaces compared with 25.7% of LOF and 9.5% of ClinVar non-pathogenic mutations, with 24.6% occurring at homomeric interfaces compared with 16.9% and 7.9%, respectively (Fig 2e). The functional importance of interactions with regulatory partners is illustrated by the oncogenic KRAS Q61H substitution, which impairs GAP-mediated GTP hydrolysis while retaining high-affinity interaction with RAF, enabling persistent downstream signalling [34, 35].

### LOF mutations preferentially disrupt regulatory post-translational modification

Both GOF and LOF mutations were enriched in the proximity of phosphorylation, ubiquitylation and acetylation sites relative to ClinVar non-pathogenic variants, demonstrating that cancer driver mutations are more frequently observed at post-translationally modified regulatory residues involved in protein function. GOF mutations at phosphorylation sites also showed enrichment within the surrounding ±3 residue region (30.0%) compared with both LOF (22.5%) and ClinVar non-pathogenic (18%) variants (Fig 3), suggesting that activating mutations may alter the local sequence environment surrounding phosphorylation sites rather than the phospho-acceptor residue itself. Because kinase recognition motifs typically extend beyond the modified residue, mutations within neighbouring residues may alter kinase recognition, substrate specificity or phosphorylation efficiency, thereby rewiring downstream signalling pathways while preserving the overall protein fold. For example, the lung cancer-associated EGFR P1019L substitution alters the sequence motif surrounding phosphorylated Tyr1016 from pYxxP to pYxxL, switching recruitment from a CRK-associated complex to a SHIP2-associated complex and promoting sustained ERK and AKT signalling, cell migration and invasion [36].

**Figure 3.**
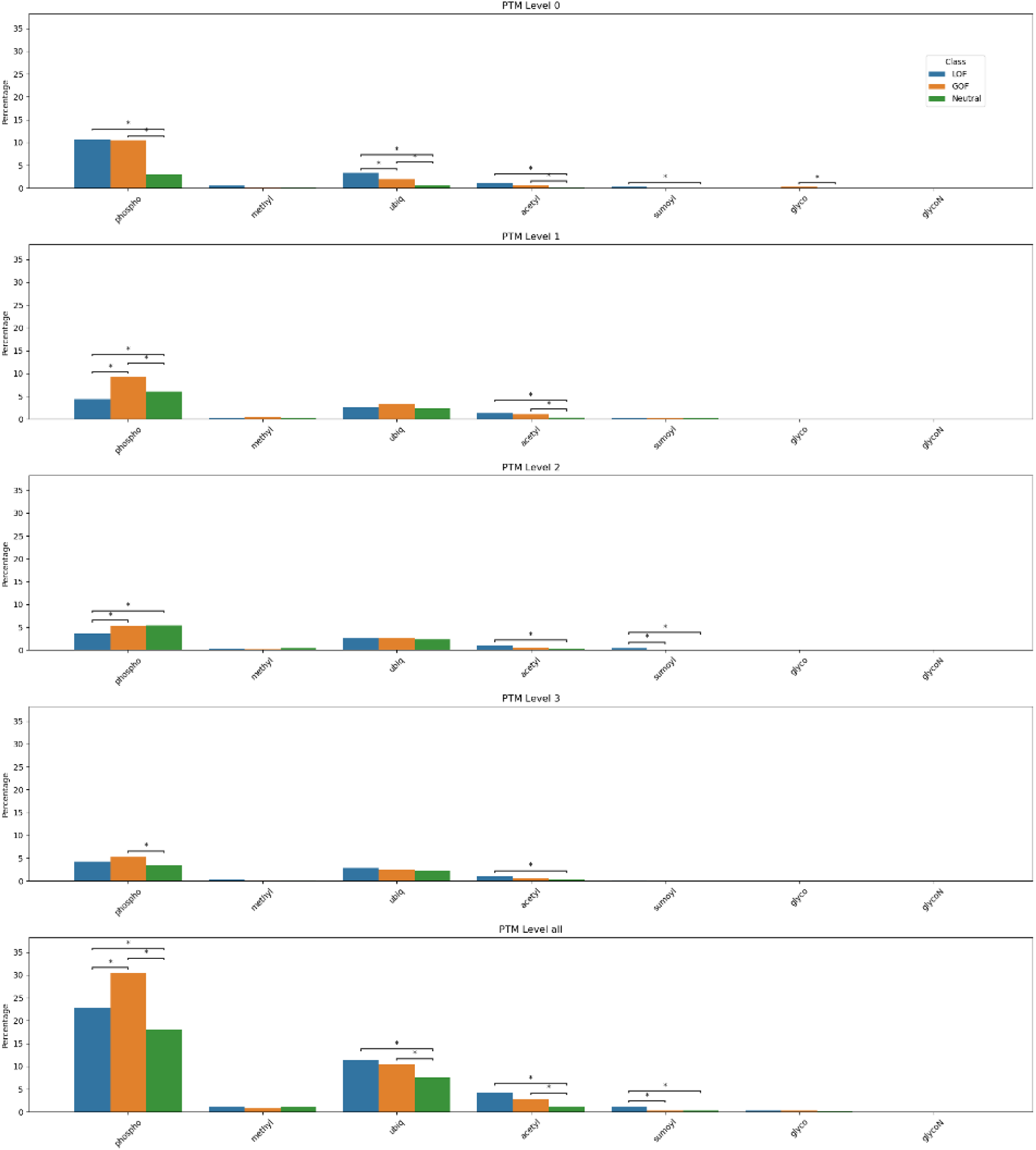
Distribution and characteristics of cancer-associated somatic mutations. (Across all panels, bar plots show the percentage of Loss-of-Function (LOF), Gain-of-Function (GOF), and ClinVar non-pathogenic mutations. Brackets with asterisks denote statistically significant pairwise differences using Fisher’s exact test). Post-translational modification (PTM) sites: Mutation frequencies across seven PTM types, stratified by distance from the target residue (Level 0: exact site; Levels 1-3: flanking positions; Level all: aggregate).

These findings suggest that GOF mutations may affect structurally important regulatory regions involved in molecular recognition and signalling rather than causing widespread disruption of protein structure.

In contrast, methylation and glycosylation sites showed no significant enrichment in proximity to GOF or LOF mutations. However, although the sample size was low, SUMOylation exhibited a distinct LOF-specific pattern with LOF mutations significantly enriched both at SUMOylated residues (0.35% versus 0.06% GOF and 0.03% ClinVar non-pathogenic) and within the surrounding ±3 residue region (1.22% versus 0.29% GOF and 0.26% ClinVar non-pathogenic) (Fig 4). Given the established roles of SUMOylation in regulating protein stability, DNA repair and cell-cycle control, these findings suggest that disruption of SUMO-mediated regulatory mechanisms represents an additional pathway contributing to protein inactivation.

**Figure 4.**
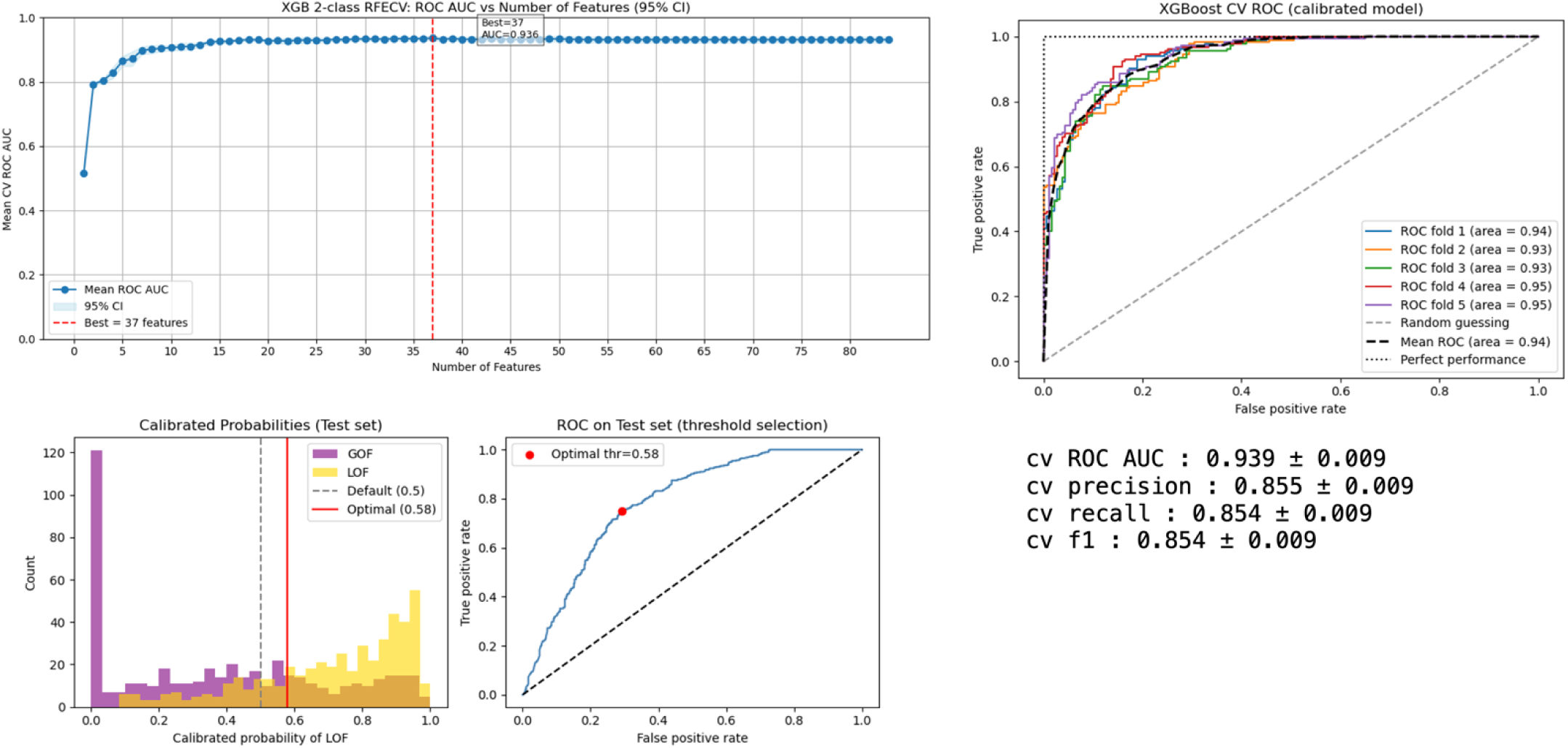
5-fold cross-validation ROC curves for the calibrated MOKCa-3D-GL, XGBoost model. The classifier achieved a mean AUC of 0.94 (black dashed line), with individual fold performances ranging from 0.93 to 0.95, demonstrating stable and accurate discriminative performance relative to the random guessing baseline (Gray dashed line). The optimal feature subset contains 37 features, achieving a maximum AUC of 0.936 (red dashed line).

### Structural and functional signatures as machine learning features

The distinct structural and functional signatures identified in the preceding analyses suggested that combining complementary biological descriptors could improve discrimination between gain-of-function (GOF) and loss-of-function (LOF) mutations. To capture higher-order relationships that are not represented by individual annotations alone, we generated 25 manually engineered composite features describing protein structural environment, physicochemical perturbation, structural constraint and non-linear interactions between complementary descriptors.

### Machine learning distinguishes pathogenicity and mutation mechanism

As an initial benchmark, we first evaluated whether these features improved conventional pathogenicity prediction by distinguishing pathogenic (GOF and LOF) mutations from ClinVar non-pathogenic variants. A Random Forest classifier achieved excellent performance, with a mean cross-validation AUC of 0.947 ± 0.033, with F1 score of 0.881 for both pathogenic and ClinVar non-pathogenic, on unseen protein variants in test and an external validation AUC of 0.8176, with F1 score 0.69 (ClinVar non-pathogenic) and 0.79 (pathogenic). This compared favourably to using AlphaMissense pathogenicity scores alone (AUC. F1), although prediction was driven primarily by the AlphaMissense pathogenicity score together with structural features. Because pathogenicity prediction is a well-established task, detailed model evaluation [28], feature importance analysis and validation results are presented in the Supplementary Results (Supplementary Fig S2–S3, Supplementary Table S2).

Next, we investigated the more challenging task of discriminating GOF from LOF mutations. Recursive Feature Elimination with Cross-Validation (RFECV) identified an optimal subset of 37 features for a XGBoost classifier (MOKCa-3D-GL) achieving a mean cross-validation AUC of 0.939 ± 0.009 (Fig 4; Table 1, Supplementary Table S3).

**Table 1.** Mean cross-validation performance metrics (AUC, Precision, Recall, and F1-score) and their standard deviation for MOKCa-3D-GL.

| **Metric** | **Performance** | **Standard Deviation** |
|---|---|---|
| **CV AUC** | 0.936 | ± 0.009 |
| **CV Precision** | 0.855 | ± 0.009 |
| **CV Recall** | 0.854 | ± 0.009 |
| **CV F1** | 0.854 | ± 0.009 |

Overall, the MOKCa-3D-GL classifier generalised well to an independent validation dataset containing variants from both proteins that were represented in the training set and proteins that were entirely unseen during training, achieving an external validation AUC of 0.834. Using an optimised decision threshold of 0.58, MOKCa-3D-GL achieved recall values of 0.74 for LOF mutations and 0.83 for GOF mutations. Corresponding external validation F1-scores were 0.79 for GOF mutations and 0.78 for LOF m utations (Fig 5), demonstrating good discrimination.

**Figure 5.**
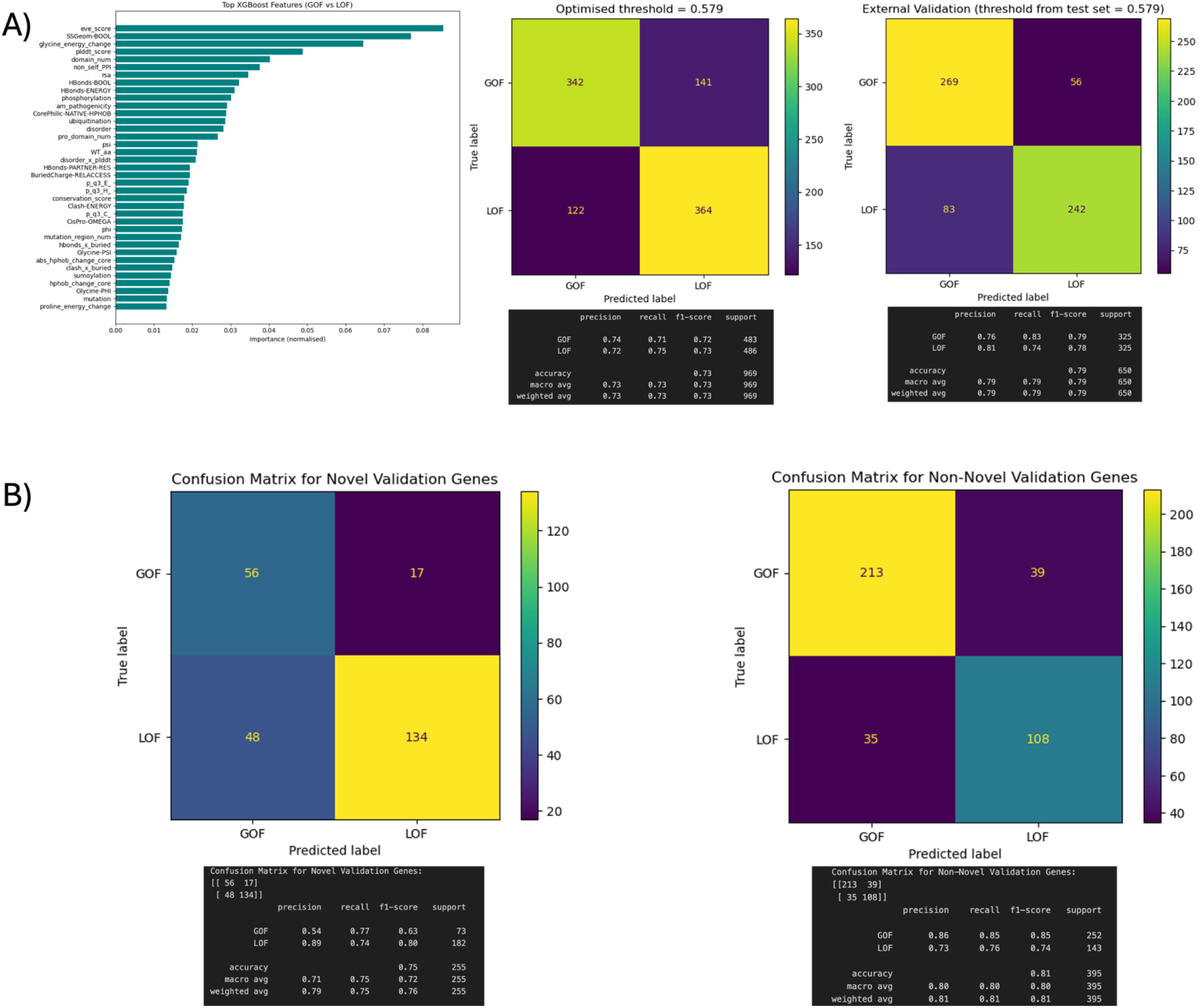
External Validation of MOKCa-3D-GL. A) Feature importance ranking, and test set, external validation set confusion matrix demonstrating model performance. B) External validation set confusion matrix demonstrating model performance for seen proteins and unseen (novel) proteins during training.

However, performance differed when stratifying the external validation set by protein novelty. For variants in previously unseen proteins, MOKCa-3D-GL achieved an F1-score of 0.80 for LOF variants but only 0.63 for GOF variants. In contrast, for variants from proteins represented during training, F1-scores were 0.74 for LOF and 0.85 for GOF (Fig 5, Supplementary Tables S4 and S5). These results suggest that LOF-associated patterns generalise more consistently to novel proteins, whereas GOF-associated patterns are less transferable across unseen protein contexts.

One possible explanation is the comparatively limited diversity of the GOF training data, which contained variants from approximately 200 proteins, compared with 334 proteins in the LOF dataset. Alternatively, GOF variants may occupy a broader and more heterogeneous feature space, making it more difficult for the model to learn generalisable patterns from the available training data.

We also trained a multi-class model (MOKCa-3D-3W; GOF, LOF & ClinVar non-pathogenic), the best performing XGB model achieved Cross Validated OvR AUC of 0.940 ± 0.006, and Macro OvR AUC 0.8889 on unsee proteins variants test set, and 0.7874 Macro OvR AUC on external validation set (shown Supplementary Fig S4-S5, Supplementary Table S6).

### 2.5 Model interpretation using SHAP

To investigate the biological basis of the classifier, model predictions were interpreted using SHAP (SHapley Additive exPlanations), which quantifies the contribution of individual features to each prediction [37] (Supplementary Fig S6). The highest-ranking features included both individual structural descriptors and several manually engineered composite features, demonstrating that mechanistic classification depended on integrating complementary structural, functional and evolutionary information rather than any single annotation.

Notably, composite features such as disorder × pLDDT, clash × buried charge, and Hbond disruption × buried charge ranked among the strongest contributors to model performance. These features capture higher-order relationships between protein flexibility, structural confidence, solvent accessibility and structural impact that are not represented by individual descriptors alone. Their importance indicates that combining complementary biological information substantially improves discrimination between GOF and LOF mutations.

Importantly, the SHAP analysis was consistent with the biological patterns identified in the preceding structural and functional analyses. Features associated with structurally constrained protein cores contributed predominantly to LOF predictions, whereas features describing regulatory domains, protein–protein interaction interfaces and local regulatory environments contributed more strongly to GOF classification. Together, these findings demonstrate that the classifier learns biologically meaningful determinants of mutation mechanism rather than relying solely on general measures of pathogenicity.

### Comparison Test

To compare the performance of the MOKCa-3D-3W multi-class, classifier with LoGoFunc, we used the DeLong test [38] using a data set of 252 GOF, 315 LOF, and 517 ClinVar non-pathogenic mutations. Although these mutations were not in our training set, the training set for LoGoFunc is confidential, so it is possible that these mutations contributed to the training of LoGoGFunc particularly for GOF mutations which are enriched in cancers. Despite this, cancer-specific training appears to confer a substantial advantage for identifying LOF variants. For direct GOF versus LOF discrimination, MOKCa-3D-GL showed a modest numerical advantage (Fig 6 and Table 2).

**Figure 6.**
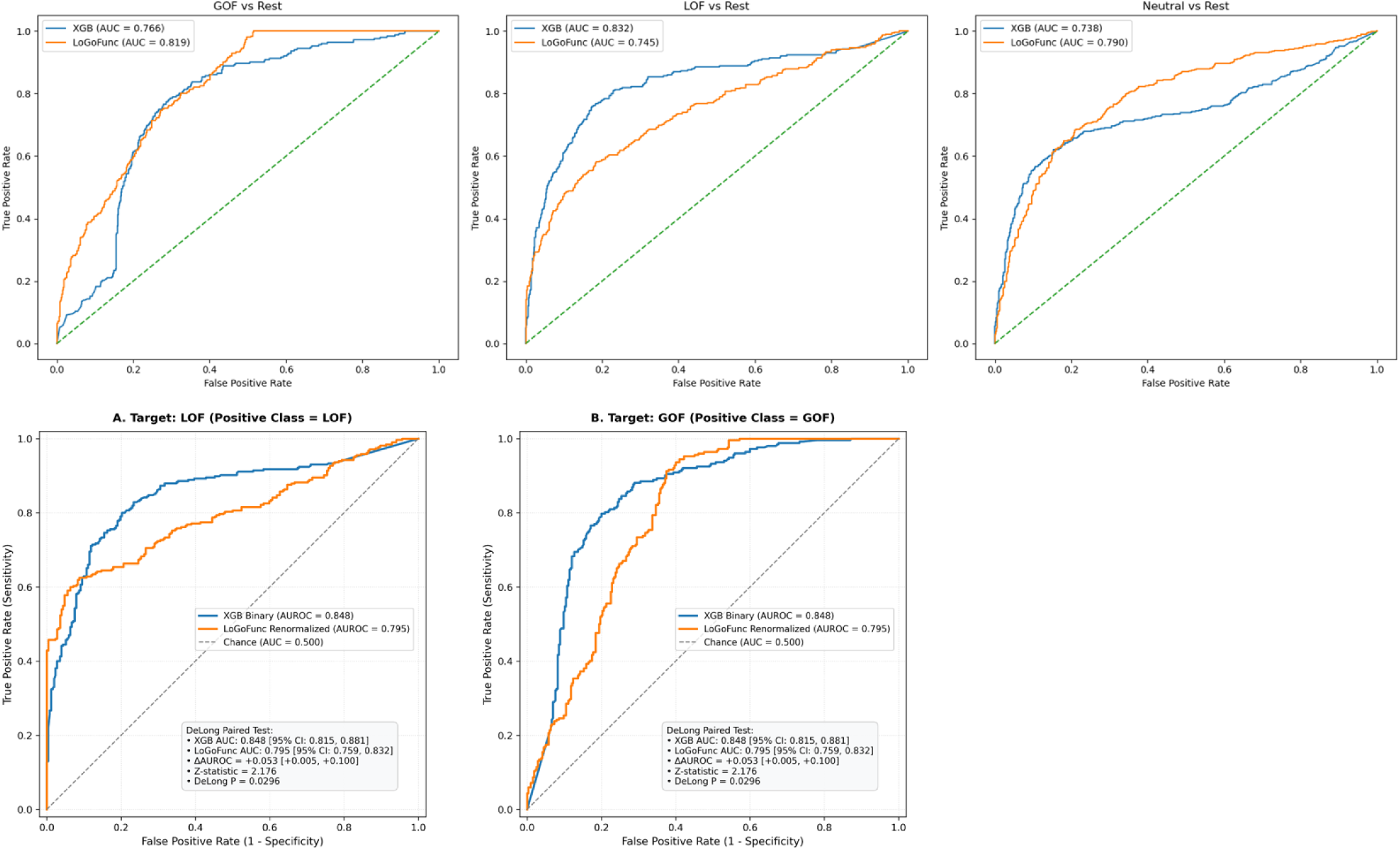
A) Multi-class Comparison of MOKCa-3D-3W models with LoGoFunc using DeLong test. B) GOF versus LOF Comparison of MOKCa-3D-GL (XGB) models with LoGoFunc using de long test.

**Table 2.** De long test results.

| **Comparison** | **MOKCa-3D**<br>**AUROC** | **LoGoFunc**<br>**AUROC** | **Delta**<br>**AUROC** | **Holm-adjusted P** |
|---|---|---|---|---|
| **GOF vs Rest** | 0.7659 | 0.8189 | -0.0530 | 0.0050 |
| **LOF vs Rest** | 0.8320 | 0.7453 | 0.0867 | 0.0001 |
| **Neutral vs Rest** | 0.7375 | 0.7902 | -0.0527 | 0.0050 |
| **GOF vs LOF** | 0.848 | 0.795 | 0.053 | 0.0296 |

### MOKCa-3D

The curated GOF and LOF annotations generated in this study and the GOF/LOF predictions have been incorporated into the MOKCa-3D database [29], providing a publicly available resource that links experimentally validated mechanistic annotations to structural and functional protein features for future studies of cancer-associated missense mutations.

## Discussion

Accurate discrimination of gain-of-function (GOF) and loss-of-function (LOF) missense mutations remains a major challenge in cancer genomics because both classes promote tumorigenesis, but through distinct molecular mechanisms. Knowledge of whether a driver mutation is GOF or LOF dictates the therapeutic approach in treating a cancer, with GOFs often being directly addressable by drugs that inactivate or inhibit the affected protein, while LOF mutations must be addressed indirectly through therapeutic targeting of a second protein whose inhibition or inactivation is synthetically lethal with the LOF-mutated protein [39]. By integrating experimentally curated mechanistic annotations with structural, functional and evolutionary protein features, we demonstrate that GOF and LOF mutations occupy distinct structural and functional environments that enable accurate mechanistic prediction across diverse cancer-associated proteins. Thus, while pathogenicity predictors identify functionally important variants, discrimination between activating and inactivating mutations requires additional structural and mechanistic information.

LOF mutations were consistently associated with features indicative of structural destabilisation, such as hydrogen bond disruption, steric clashes and buried charge substitutions, supporting protein inactivation through disruption of the native fold. In contrast, GOF mutations preferentially occurred at solvent-accessible residues and exhibited fewer destabilising features, suggesting that oncogenic activation is more commonly achieved through subtle modulation of protein regulation and molecular interactions than through widespread structural disruption.

Structural features alone did not fully distinguish GOF from LOF mutations. Functional annotations revealed additional differences: GOF mutations were enriched at protein–protein interaction interfaces and near phosphorylation sites, whereas LOF mutations were enriched near SUMOylation sites. These findings suggest that mutation mechanism is reflected across multiple levels of protein organisation, from local structural stability to higher-order regulatory and signalling networks, explaining why integration of complementary structural and functional features substantially improved mechanistic classification.

Our findings extend previous studies of mutation mechanism. Gerasimavicius et al. [21] demonstrated, using predominantly inherited disease mutations, that GOF, LOF and dominant-negative variants occupy distinct structural environments, while Singh et al. [22] showed that activating and inactivating mutations in protein kinases can be distinguished using structural and evolutionary features. We demonstrate that these principles extend to experimentally characterised somatic cancer mutations across diverse protein families and further show that functional protein annotations including interaction interfaces, protein domains and post-translational modification sites provide complementary information beyond structural descriptors alone.

Although these observations broadly agree with studies of inherited disease, somatic cancer mutations evolve under fundamentally different selective pressures. Germline pathogenic variants are selected according to organismal fitness, whereas somatic driver mutations are selected for their effects on tumour cell fitness. Consequently, cancer-associated GOF mutations may exploit functional alterations that are tolerated during protein evolution but become oncogenic by conferring constitutive activity, altered substrate specificity or novel molecular interactions. Similarly, cancer-associated LOF mutations can arise through disruption of regulatory interactions or domain-specific functions in addition to classical structural destabilisation. These differences emphasise the need for cancer-specific mechanistic prediction methods.

The biological differences identified in this study translated naturally into predictive performance. By integrating structural, functional and engineered descriptors, we developed a structure-informed machine learning framework capable of distinguishing GOF and LOF mutations with high accuracy. Unlike existing sequence-based predictors such as AlphaMissense, which primarily distinguish deleterious or pathogenic variants from tolerated variation, our classifier was designed specifically to predict molecular mechanism. Engineered composite features contributed substantially to model performance, demonstrating that biologically informed feature engineering captures higher-order relationships beyond individual annotations.

Although our curated dataset represents one of the largest collections of experimentally validated cancer GOF and LOF missense mutations currently available, some protein families remain underrepresented and mechanistic annotations remain limited by the available experimental literature. Continued experimental characterisation together with community-driven curation will therefore be essential for developing larger benchmark datasets and improving future mechanistic predictors.

In addition to the predictive framework, this work provides a community resource. The curated GOF and LOF annotations and corresponding predictions have been incorporated into the MOKCa-3D database [29], where each mutation is linked to supporting experimental evidence together with structural and functional annotations. This resource provides a benchmark for future development and evaluation of computational methods for mechanistic interpretation of cancer driver mutations.

In summary, our study demonstrates that GOF and LOF mutations occupy distinct structural and functional environments that can be exploited for accurate mechanistic prediction across diverse cancer-associated proteins. More broadly, these findings show that integrating complementary structural and functional protein annotations enables prediction of mutation mechanism beyond conventional pathogenicity assessment, providing a foundation for improved interpretation of somatic cancer variant.

Together, these findings provide new biological insights into cancer driver mutations while establishing a framework for mechanism-aware variant interpretation.

By extending observations from inherited disease to somatic cancer and integrating structural, functional and engineered protein features within an interpretable machine-learning framework, this work provides a foundation for more accurate mechanistic interpretation of cancer-associated missense mutations.

## MATERIALS AND METHODS

### Datasets

The data collection and processing step was crucial to standardize data and ensure its quality, and the overall workflow is shown in Supplementary Fig S7. For the training and test set, gain-of-function (GOF) and loss-of-function (LOF) cancer-associated missense mutations were manually curated from the primary literature. Non-pathogenic mutations were extracted from ClinVar [30] and were used as the ClinVar non-pathogenic dataset. The final dataset comprised 1,727 GOF mutations from 200 proteins, 2,857 LOF mutations from 336 proteins, and 3831 ClinVar non-pathogenic variants (334 proteins). The validation set of mutations were obtained from OncoKB, comprising 325 GOF mutations (22 proteins), and 325 LOF mutations (20 proteins) and 550 ClinVar non-pathogenic mutations, none of which were present in the original datasets.

### Structural Annotation

The structural impact of each mutation was characterised using the SAAP-MOKCa-3D pipeline [29]. SAAP models each amino acid substitution by generating a local structural model from the corresponding AlphaFold-predicted protein structure and calculates fourteen sequence- and structure-based descriptors that characterise the predicted structural consequences of the variant [40, 41]. These descriptors include hydrogen bond disruption, disulphide bond disruption, steric clashes, cavity formation, ligand-binding residues, electrostatic charge changes, hydrophobicity changes, residue conservation, and UniProtKB/Swiss-Prot functional annotations. Residue-level pLDDT confidence scores were extracted from AlphaFold [42] Protein Structure Database models and included as a measure of local structural confidence.

NetSurfP3 was also used to predict wild-type residue structural properties, including relative solvent accessibility (RSA), absolute solvent accessibility (ASA), secondary structure (helix, strand and coil) and intrinsic disorder [43]. Residues with an RSA ≤25% were classified as buried, and residues with a disorder probability >0.5 were classified as disordered.

### Functional Annotation

Functional annotations were generated using established computational predictors and publicly available biological databases.

Protein–protein interaction interface residues were annotated using PIONEER2.0 a deep learning framework that predicts protein-binding partner-specific interfaces for known protein interactions across the human proteome [44]. Each mutation was assessed based on whether the PIONEER-predicted interface residue was involved in participated in homomeric (self) or heteromeric (protein–protein) interaction interfaces.

Pfam protein domain annotations were extracted from InterPro [45] to determine if a mutation was located within a protein domain.

Post-translational modification (PTM) annotations were obtained from PhosphoSitePlus [46], a comprehensive knowledgebase for mammalian PTMs, including phosphorylation, acetylation, ubiquitylation, Glycosylation, SUMOylation, and methylation. Mutations were classified as PTM-associated if they occurred at the modified residue or within three residues upstream or downstream of a documented PTM site.

### Pathogenicity Annotation

The AlphaMissense pathogenicity score was included as one of the predictive features AlphaMissense [14] is a deep learning–based predictor of missense variant pathogenicity that integrates protein language model representations with structural information derived from AlphaFold [14]. Although it achieves state-of-the-art performance for distinguishing pathogenic from benign variants, it was not designed to discriminate between GOF and LOF mutations.

Evolutionary model of variant Effect (EVE) [12] is a model for the prediction of clinical significance of human variants based on sequences of diverse organisms across evolution. It uses fully unsupervised deep learning trained on amino acid sequences of over 140K species. An EVE pathogenicity score was included as a predictive feature, which range from 0 to 1 (0 represent the most benign/neutral variant and 1 represent the most pathogenic variant). Evolutionary Scale Modeling (ESM) [13, 18] protein language models, estimate the functional impact or likelihood of amino acid mutations in a protein sequence. They assign log-probability or likelihood scores to specific residues or substitutions based on evolutionary patterns learned from millions of protein sequences. Higher or positive values suggest enhanced or tolerated fitness, while strongly negative values indicate that a mutation is disruptive, damaging, or functionally constrained.

### Statistical Analysis

Pairwise statistical comparisons were performed between gain-of-function (GOF), loss-of-function (LOF) and ClinVar non-pathogenic variants to identify differences in structural, functional and pathogenicity features. Continuous variables were categorised using predefined thresholds prior to analysis. Fisher’s exact test, implemented in the SciPy python library [47] was used to compare categorical variables. To account for multiple testing, *P*-values were adjusted using the Bonferroni correction.

### Feature Engineering

To improve predictive performance, individual structural, functional, pathogenicity and sequence-derived annotations were integrated into a unified feature set for machine learning.

### Feature Categories

The final feature set comprised four complementary categories of descriptors. Structural features included SAAP-derived descriptors, relative and absolute solvent accessibility, secondary structure, intrinsic disorder and residue-level pLDDT confidence scores. Functional features comprised protein–protein interaction interfaces, Pfam domain annotations and post-translational modification sites. Pathogenicity features consisted of the AlphaMissense pathogenicity score and EVE score, while sequence-derived features included residue conservation score, ESM score (ESM-1b) obtained from ProtVar database [18] and additional sequence-based descriptors generated by the SAAP pipeline.

### Composite Feature Engineering

In addition to experimentally derived and computationally predicted annotations, we manually designed 25 biologically motivated composite features to capture higher-order structural relationships that are not represented by individual descriptors alone. These engineered features were developed using structural biology domain knowledge to improve class separability and represented four principal biological themes: (i) protein structural environment, combining solvent accessibility with local structural rigidity to distinguish solvent-exposed loop regions from buried protein cores; (ii) physicochemical perturbation, quantifying the magnitude of changes in residue charge and hydrophobicity; (iii) structural constraint, identifying mutations affecting structurally critical glycine and proline residues; and (iv) non-linear feature interactions, combining complementary descriptors to model complex relationships between structural properties.

### Model Development

Three classification datasets were constructed to address complementary prediction tasks. **(i)** A binary pathogenicity dataset (Pathogenic vs. ClinVar non-pathogenic) comprised 2,800 training variants per class and an independent external validation set of 550 neutral variants and 650 pathogenic variants (which have equal 325 GOF and LOF variants). (ii) A three-class dataset (GOF, LOF and ClinVar non-pathogenic) comprised 1,400 training variants per class and 325 external validation variants per class (iii) A binary mechanistic dataset (GOF vs. LOF) comprised 1,400 training variants per class and an independent external validation set of 325 variants per class. Based on the characteristics of the feature set and their established performance in structured biological datasets, three ensemble machine learning algorithms were evaluated: Random Forest, XGBoost and LightGBM. Models were trained using stratified five-fold cross-validation to preserve class distributions. A fixed random seed (random_state = 42) was used to ensure reproducibility. Gridsearch hyperparameters tuning was applied (Supplementary Fig. S8). All code and data are available in GitHub: https://github.com/UOSbioinformaticslab/MOKCa-GofLof.

### Recursive feature elimination

To identify the most informative predictive features and reduce model complexity, Recursive Feature Elimination with Cross-Validation (RFECV) was performed independently for each classification task. The optimal feature subset identified by RFECV was subsequently used to train the final classification model.

### Model evaluation

Model performance was assessed using the area under the receiver operating characteristic curve (AUC) and F1-score, with the highest-performing model selected for subsequent analyses. Feature importance was quantified using SHAP (SHapley Additive exPlanations) values calculated using the TreeSHAP [37] algorithm implemented in the SHAP Python package. Global feature importance was estimated using the mean absolute SHAP value across all validation samples.

### Model Validation

Two set of mutations were obtained (i) other mutations in the proteins used to train the classifier (252 GOF, and 143 LOF) (ii) mutations from proteins that are not in the data set (73 GOF, and 182 LOF).

### Comparison with LoGoFunc model

Model discrimination was compared using the nonparametric paired DeLong test for correlated ROC curves, as both MOKCa-3D-GL XGB model and LoGoFunc generated predictions for the same external validation variants. AUROC was calculated using continuous prediction probabilities, and the difference between correlated AUROCs was tested using the DeLong variance-covariance estimator with a two-sided test. For the three-class task, one-versus-rest AUROCs were calculated separately for GOF, LOF, and ClinVar non-pathogenic variants predicted using multi-class MOKCa-3D-3W XGB model and LoGoFunc. For direct GOF-versus-LOF discrimination, ClinVar non-pathogenic variants were excluded. Because LoGoFunc provides three-class probabilities, its GOF score for this analysis was converted to a conditional probability, 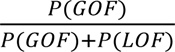, and LOF scores converted to conditional probability, 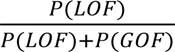, whereas the binary MOKCa-3D-GL XGB GOF probability was used directly. The three one-versus-rest P-values were corrected using the Holm–Bonferroni procedure, with adjusted P < 0.05 considered statistically significant. AUROC differences were reported as XGBoost minus LoGoFunc.

7.

## Abbreviations

GOF: Gain-of-function
LOF: Loss-of-function
XGB: Extreme Gradient Boosting
LightGBM: Light Gradient Boosting Machine
CV: Cross Validation
AUC: Area Under the Receiver Operating Characteristic Curve
vs: Versus

## Ethics approval and consent to participate

Not applicable. Computational analysis of previously reported cancer associated variants.

## Consent for publication

Not applicable.

## Availability of data and materials

All data used in this study were obtained from publicly available sources. GOF and LOF cancer-associated missense mutations in the training and test sets were manually curated from the primary literature, and the PubMed IDs to the assignment recorded. The mutations have are incorporated into the MOKCa-3D database (https://bioinformaticslab.sussex.ac.uk/MOKCa-3D/) alongside non-functionally assigned cancer-associated missense mutations.

Mutations described as non-pathogenic in the ClinVar database were used as the set of neutral mutations. Mutations used to validate the ML methods were extracted from OncoKB.

All analysis code used in this study is available in the GitHub repository: https://github.com/UOSbioinformaticslab/MOKCa-GofLof

## Authors’ contributions

Contribution: BH developed the software, analysed the data, designed figures and wrote the manuscript; FP designed the research, analysed the data, supervised the study and wrote the manuscript.

## Acknowledgements

We would like to thank Laurence Pearl for critical reading of the manuscript.

